# Programming T cells for intercellular genome editing

**DOI:** 10.64898/2026.06.21.729417

**Authors:** Kevin M. Wasko, Madeleine Maker, Wayne Ngo, Kai Chen, Enbo Ma, Rithu Pattali, Enzo Chen, Trevor Leung, Jonathan Braverman, Jennifer A. Doudna

## Abstract

Therapeutic genome editing requires delivery of editing molecules to defined cell types, but targeting specificity and efficiency are currently limited. We hypothesized that properties inherent to immune cells, including tissue infiltration and programmed cell recognition, could be harnessed to engineer a cell-based delivery system. We show here that T cells can both produce and transfer editing machinery to target cells. In response to a programmable ligand, engineered T-lymphoid cells can transfer enzymes using complex spatiotemporal logic and deliver cargo in a cell contact-dependent or -independent manner. We demonstrate feasibility of this approach in primary human T cells, establishing a customizable genetic circuit for macromolecular delivery controlled by intercellular interactions.

## Introduction

Therapeutic genome editing requires precise *in vivo* delivery of editing enzymes to maximize activity in target cells while minimizing genome modification in off-target tissues. Current nonviral delivery strategies use synthetic or biologically derived particles to transport editing enzymes into cells. When administered systemically, these carriers rely on diffusion and surface antigen binding, to reach their targets, restricting targeted delivery to cell types with unique surface markers^1^.

We previously developed enveloped delivery vehicles (EDVs), virus-inspired particles that package CRISPR-Cas9 ribonucleoproteins (RNPs) fused to a truncated HIV-1 Gag polyprotein (miniGag) for targeted cell editing^2,3^. While this approach is highly effective for inducing genome edits in human T cells *in vivo*^4,5^, the number of T cells directly transduced is small. Activation-induced clonal expansion naturally increases the number of edited T cells to a therapeutically useful population, a mechanism not available in other tissue types. Increasing the local concentration of delivery vehicles within a target tissue could overcome this limitation. Because immune cells can actively home to specific tissues and integrate signals from multiple antigens simultaneously, they are potentially well suited for targeted production and delivery of editing enzymes.

To test this idea, we engineered immune cells to possess both Cas9-EDV production capabilities and the receptor-activated behavior of T cells. We constructed a Juxtacrine Enzyme Transfer (JET) delivery circuit in which a synthetic Notch receptor (SNIPR) controls EDV production, thereby enabling packaged editor production at the site of receptor engagement. We show that JET T cells can respond to a variety of ligands by transferring molecules using either EDV-based or direct cell-cell mechanisms. JET T cells could overcome the limitations of existing delivery systems by using cellular logic to enable local production and delivery of biological cargo.

## Results

### Inducible intercellular CRISPR-Cas9 transfer by Jurkat T cells

Inspired by the transfer of protein and RNA cargo using virological synapses between CD4^+^ cells during HIV-1 replication^6–8^, we wondered whether T cells could similarly transfer cargo to neighboring cells using EDVs. In Jurkat cells, an immortalized CD4^+^ T cell line, we genomically integrated a doxycycline-inducible EDV construct, containing a miniGag-Cas9 fusion and the vesicular stomatitis virus glycoprotein (VSV-G), along with a neomycin resistance cassette for selection^9,10^ (Fig. 1A). As controls, we tested constructs that lacked either miniGag, which localizes Cas9 to the plasma membrane, or VSV-G, which mediates endosomal escape of cargo, or both (Fig. S1A). We confirmed the expression of each component in the presence of doxycycline (Fig. 1B).

**Figure 1:**
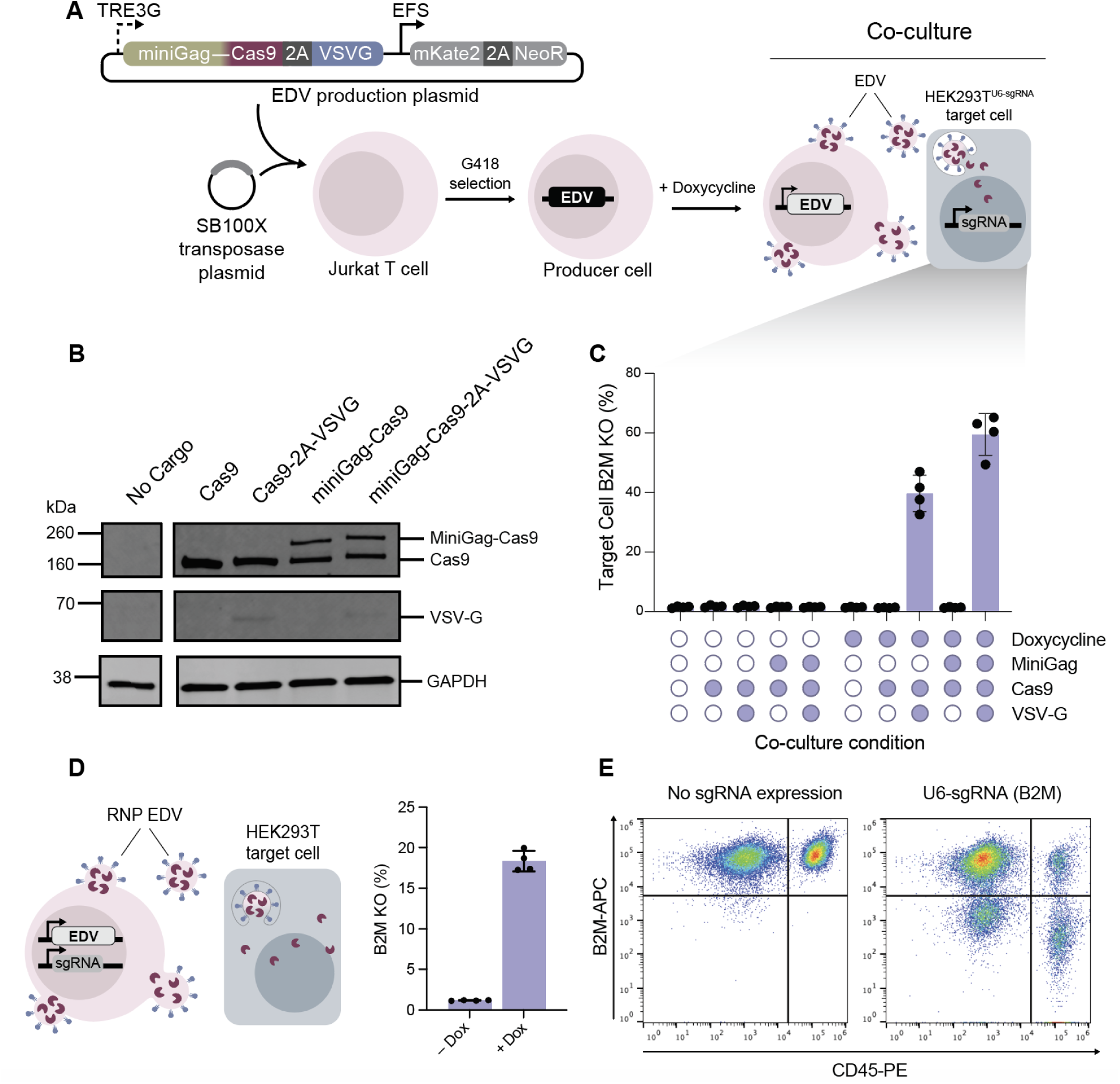
Doxycycline-inducible CRISPR-Cas9 transfer by Jurkat T cells. A. Plasmid map and producer Jurkat cell production schematic. Jurkats are nucleofected with Sleeping Beauty plasmids encoding inducible Cas9 EDV cassettes, selected with the antibiotic G418, and co-cultured with target cells harboring a sgRNA. B. Western blots confirm expression of FLAG-tagged Cas9 or miniGag-Cas9 and VSV-G in Jurkat cells cultured with doxycycline. C. *B2M* knockout measured by flow cytometry in HEK293T target cells following 6 days of co-culture with producer Jurkats in the presence or absence of doxycycline. (*n* = 4 biological replicates) D. *B2M* knockout indicating Cas9 RNP transfer following co-culture of unmodified HEK293T cells and producer cells with genomically integrated EDV and enU6-sgRNA cassettes. (*n* = 4 biological replicates) E. Representative flow cytometry plot displaying *B2M* knockout in both producer (CD45^+^) and target (CD45^−^) cells.

We first tested whether Cas9 protein can be transferred in the absence of a single guide RNA (sgRNA). To monitor functional Cas9 apoprotein delivery, the engineered Jurkat cells were co-cultured with HEK293T cells expressing a sgRNA targeting the endogenous beta-2 microglobulin gene locus. We observed loss of β2M protein expression in target cells that depended on both doxycycline and EDV components, indicating successful Cas9 transfer (Fig. 1C). No β2M loss was observed upon expression of Cas9 or miniGag-Cas9 alone, ruling out fusogen-independent mechanisms (Fig. 1C). Knockout in the absence of miniGag occurred with lower efficiency, consistent with VSV-G’s ability to drive budding of vesicles that could nonspecifically package cytosolic Cas9^11^.

To investigate whether T cells can also produce and transfer functional Cas9-sgRNA ribonucleoproteins (RNPs), we generated producer cells expressing both Cas9 EDVs and *B2M*-targeting sgRNA. We found that a CMV enhancer upstream of the U6 promoter driving sgRNA expression improved the rates of RNP transfer by at least two-fold^12,13^ (Fig. S1B-D). Using this enhancer-U6 construct, we observed up to 18% β2M loss in unmodified HEK293T cells following co-culture (Fig. 1D, E), demonstrating that engineered T lymphoid cells can be induced to produce and deliver fully functional genome editors to neighboring cells.

### Jurkat T cells produce EDVs for primary cell delivery

Reasoning that EDV designs optimized for production in HEK293T cells may not be optimal in other cell types, we tested additional fusogens and scaffold assemblies. Replacing VSV-G with Cocal-G, a fusogen resistant to inactivation by human serum^14^, mediated equivalent Cas9 delivery in co-culture (Fig. S2A, B). We next replaced the EDV structural scaffold, miniGag, with an enveloped protein nanocage (EPN24), a chimeric protein engineered to assemble into a 60-subunit dodecahedron, which has previously been fused to RNA binding proteins to mediate intercellular transfer of mRNA^15–17^. EPN24 did not improve Cas9 transfer efficiency to co-cultured cells relative to constructs lacking a membrane localization domain altogether (Fig. S2C, D). We reasoned that the hyperstability of the quaternary structure of EPN24 could hinder Cas9 function in target cells, so we created an alternate version of EPN that trimerizes but does not form 60-subunit cages^18^ (Fig. 2A). Unexpectedly, this trimeric construct outperformed both EPN24 and miniGag in Cas9 delivery, suggesting that higher-order assembly is dispensable for efficient cargo transfer (Fig. 2B; Fig. S2D). For simplicity, we refer to this construct, which retains the membrane association and budding domains of EPN24, as the enveloped protein trimer (EPT).

**Figure 2:**
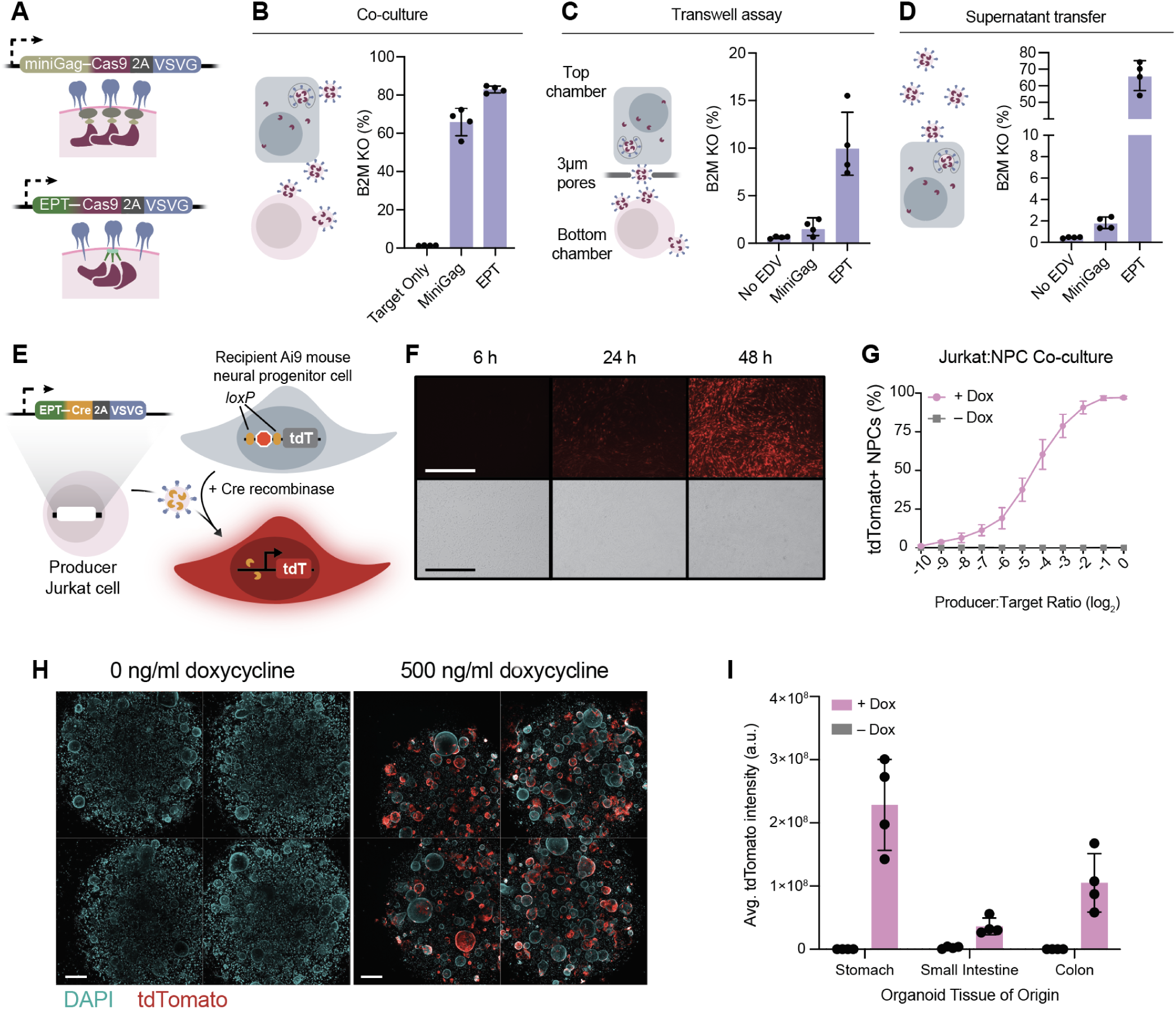
T lymphoid cells produce EDVs for primary cell delivery. A. MiniGag and EPT EDV visual representations. B-D. *B2M* knockout assessed by flow cytometry measuring transfer of Cas9 from Jurkats induced to produce miniGag- or EPT-based EDVs with doxycycline, following (B) direct co-culture, (C) co-culture in transwell assay, or (D) treatment of target cells with filtered Jurkat supernatant. (*n* = 4 biological replicates) E. Transfer of Cre results in induction of tdTomato expression in Ai9 mouse cells. F. Representative microscopy image of NPC tdTomato fluorescence in co-culture. (Producer:Target = 0.125, scale bar = 400µm) G. Flow cytometry analysis quantification of tdTomato expression in NPCs across a range of P:T ratios. (*n* = 4 biological replicates) I. Microscopy images of producer cells in co-culture with Ai9 mouse stomach-derived organoids in the presence or absence of doxycycline. Images from four wells are shown for each condition. (scale bar = 500 µm) J. Comparison of tdTomato fluorescence following 3D co-cultures +/− 500ng/ml doxycycline for organoids derived from various Ai9 mouse organs. (*n* = 4 biological replicates)

We next wanted to better understand the mechanisms of Cas9 transfer by Jurkat cells using either the miniGag or the EPT designs. We first examined whether cell-cell contact was necessary using transwell assays, in which a porous membrane (3 µm) separates producer cells from target cells but permits vesicle diffusion. In this experiment, target cell editing was more efficient for the EPT-based construct (Fig. 2C), suggesting that EPT designs generate vesicles more efficiently in Jurkat cells than miniGag. To confirm these results, we transferred filtered supernatants from Jurkat cells cultured in the presence of doxycycline to HEK293T target cells. Here too, conditioned supernatant from Jurkats engineered with EPT-based EDV cassettes mediated significantly higher editing (Fig. 2D), suggesting that while both EPT and miniGag can bud vesicles in Jurkats, EPT-based scaffolds are more efficient. Most of the cargo transfer observed for miniGag-based EDVs in Jurkats in direct co-culture likely used an intercellular contact-dependent mechanism^19^ (Fig. 1C, 2B). However, we found both constructs to bud EDVs efficiently from HEK293T producer cells (Fig. S2E, F). These results suggest that transfer mechanisms likely involve both direct membrane exchange and particle production, with different producer cell types employing each to different degrees.

To test whether this budding-optimized EPT construct is capable of efficient cargo transfer to primary cells, we utilized cells derived from *loxP*-*stop-loxP-*tdTomato (*lsl-*tdTomato) Ai9 reporter mice, which provide a fluorescent readout of Cre recombinase-mediated genome modification. We generated Jurkat cells with inducible EPT-Cre-EDV expression cassettes and performed co-culture assays with Ai9 neural progenitor cells (NPCs) at a range of producer:target (P:T) ratios (Fig. 2E). tdTomato fluorescence was visible after 24 hours of co-culture, suggesting rapid cargo transfer after genetic induction with doxycycline (Fig. 2F). Virtually all NPCs expressed tdTomato after 3 days at a P:T of 1:1, with tdTomato^+^ cells observed down to a P:T of 0.00098:1, indicative of extremely efficient Cre transfer (Fig. 2G). Transfer of supernatant from producer Jurkats to NPCs also confirmed the production of functional Cre-EDVs (Fig. S2G).

Encouraged by the efficient delivery to primary cells in a monolayer co-culture, we sought to test cargo transfer in a more physiologically relevant system that recapitulates *in vivo* tissue architecture. We performed 3D co-cultures in matrigel using wildtype organoids derived from Ai9 mouse colon, stomach, or small intestine cells. We observed tdTomato^+^ organoid cells only in the presence of doxycycline for all tissue types tested, with up to 6 or 8 percent of organoid cells edited in matrigel for small intestine and stomach organoids, respectively (Fig. 2H, I; Fig. S2H). Taken together, these results suggest that engineered lymphoid cells can produce EDVs that deliver genome editing enzymes to a variety of cell types and tissues.

### Antigen-gated EDV production enables editor transfer upon target recognition

Doxycycline provides a binary switch for controlling EDV production, but does not govern the site of delivery. To address this challenge, we developed a synthetic circuit enabling cells to produce EDVs autonomously upon sensing a protein ligand on the surface of target cells. We refer to producers engineered with this system as Juxtacrine Enzyme Transfer (JET) cells. We engineered Jurkats to express an anti-CD19 synthetic Notch receptor (SNIPR) to drive EPT-Cre-EDV expression after ligand binding^20,21^ (Fig. 3A; Fig. S3A) and generated CD19^+^ and CD19^−^ *lsl*-GFP HEK293T lines to serve as reporters of antigen-gated Cre delivery (Fig. 3B). Following three days of co-culture, we observed successful Cre transfer, with GFP expression detected in 70% of CD19^+^ cells (Fig. 3C). GFP^+^ cells were not observed following co-culture with reporter cells deficient in CD19, confirming that the SNIPR system stimulated EDV production (Fig. 3C). We used this assay to compare several SNIPR variants and found that a previously described hinge mutation enabled potent cargo transfer with minimal toxicity to producer cells^21^ (Fig. S3B-C).

**Figure 3:**
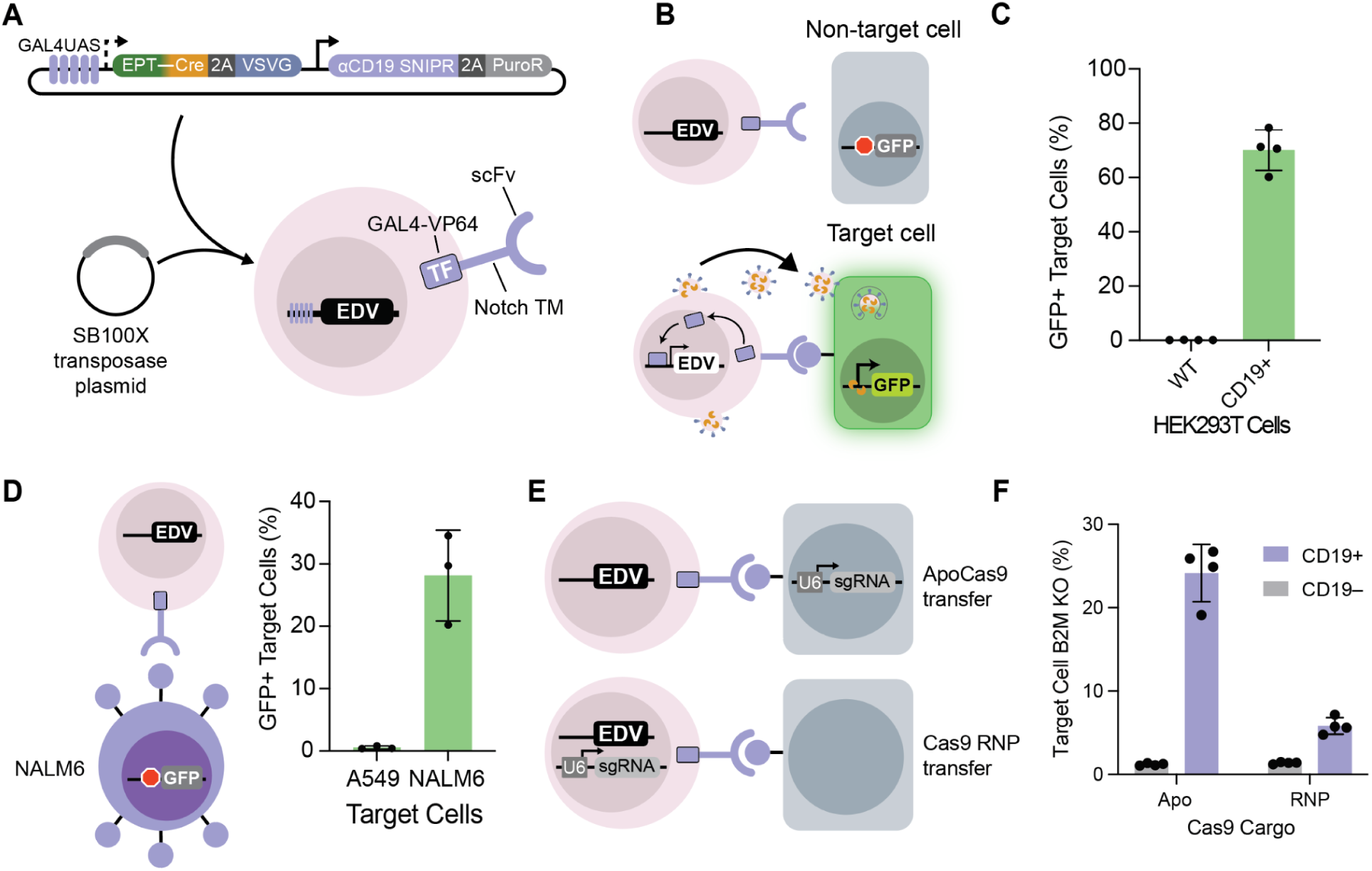
Antigen-gated EDV production enables editor transfer upon target recognition. A. Plasmid map and producer JET Jurkat production schematic. B. Engagement of the SNIPR with CD19 drives expression of Cre EDVs. Delivery of Cre to target cells removes *lsl*, resulting in constitutive GFP expression. C. Cre delivery assessed by flow cytometry measurement of GFP expression in CD19^−^(WT) or CD19^+^ target *lsl-*GFP HEK293T cells following 3 days of co-culture with producer Jurkat cells. (*n* = 4 biological replicates) D. Cre delivery to *lsl*-GFP target cell lines that do (NALM6) or do not (A549) express CD19 endogenously following 3 days of co-culture with JET Jurkat cells. (*n* = 3 biological replicates) E. Schematic of Cas9 transfer; to measure apoCas9 or Cas9 RNP transfer, target or producer cells, respectively are transduced with an enU6-sgRNA cassette. F. Flow cytometry measurement of *B2M* knockout in target HEK293T cells following 6 days of co-culture. (*n* = 3 biological replicates)

To ensure that the cargo transfer observed in these experiments was not an artifact of artificially overexpressed CD19, we also confirmed that CD19-targeted JET Jurkat cells were able to transfer cargo to *lsl-*GFP NALM6 cells, which endogenously express CD19, but not to *loxP*-GFP A549 cells lacking CD19 expression, indicating that physiological levels of target antigens are sufficient for activity (Fig. 3D).

In addition to cell surface markers, soluble ligands could extend sensing beyond specific cell types to broader tissue environments. Using JET Jurkat cells equipped with an anti-TGFβ1 SNIPR, we observed Cre transfer exclusively in the presence of exogenous TGFβ (Fig. S3D), confirming that extracellular cytokines can induce EDV production.

To test this system in a 3D culture, we generated shAPC, *Kras*^G12D^*;Trp53*^-/-^ mutant (AKP) mouse colon cancer organoids and introduced *lsl*-tdTomato, along with human CD19 at various levels of expression (Fig. S3E). Following co-culture with CD19-responsive Jurkat Cre-EDV producers, tdTomato^+^ cells were restricted to CD19^+^ organoids, and the overall fluorescence intensity of each co-culture directly corresponded to the level of CD19 expression for three of four JET Jurkat lines tested (Fig. S3F, G).

We then applied this approach to transfer Cas9 enzymes by engineering Jurkat cells with EPT-Cas9-EDV SNIPR circuits. We again monitored apoCas9 and Cas9 RNP transfer by integrating enU6::sgRNA cassettes targeting *B2M* in target and producer cells, respectively (Fig. 3E). Following co-culture, β2M loss was observed only in CD19^+^ targets, confirming functional delivery of both Cas9 protein and RNP (Fig. 3F). These experiments collectively demonstrate that cells can be engineered to transfer genome editors to neighboring cells via EDV production in response to contact with a programmable protein ligand.

### Delivery specificity by JET Jurkat cells is tunable at multiple circuit nodes

A central advantage of a cell-based chassis for EDV production is the ability to perform complex, context-dependent delivery. In principle, binding moiety of the SNIPR confers the specificity of editor transfer in our system. To test editor transfer modularity, we generated *lsl*-GFP K562 target cells with a panel of ectopically expressed ligands. JET Jurkat cells expressing SNIPRs with scFvs targeting each of these ligands or TGFβ were co-cultured with each K562 line combinatorially. Cre delivery was specific across all 6 scFv-ligand pairs tested, demonstrating that SNIPR programmability extends to antigen-specific control of EDV production (Fig. 4A).

**Figure 4:**
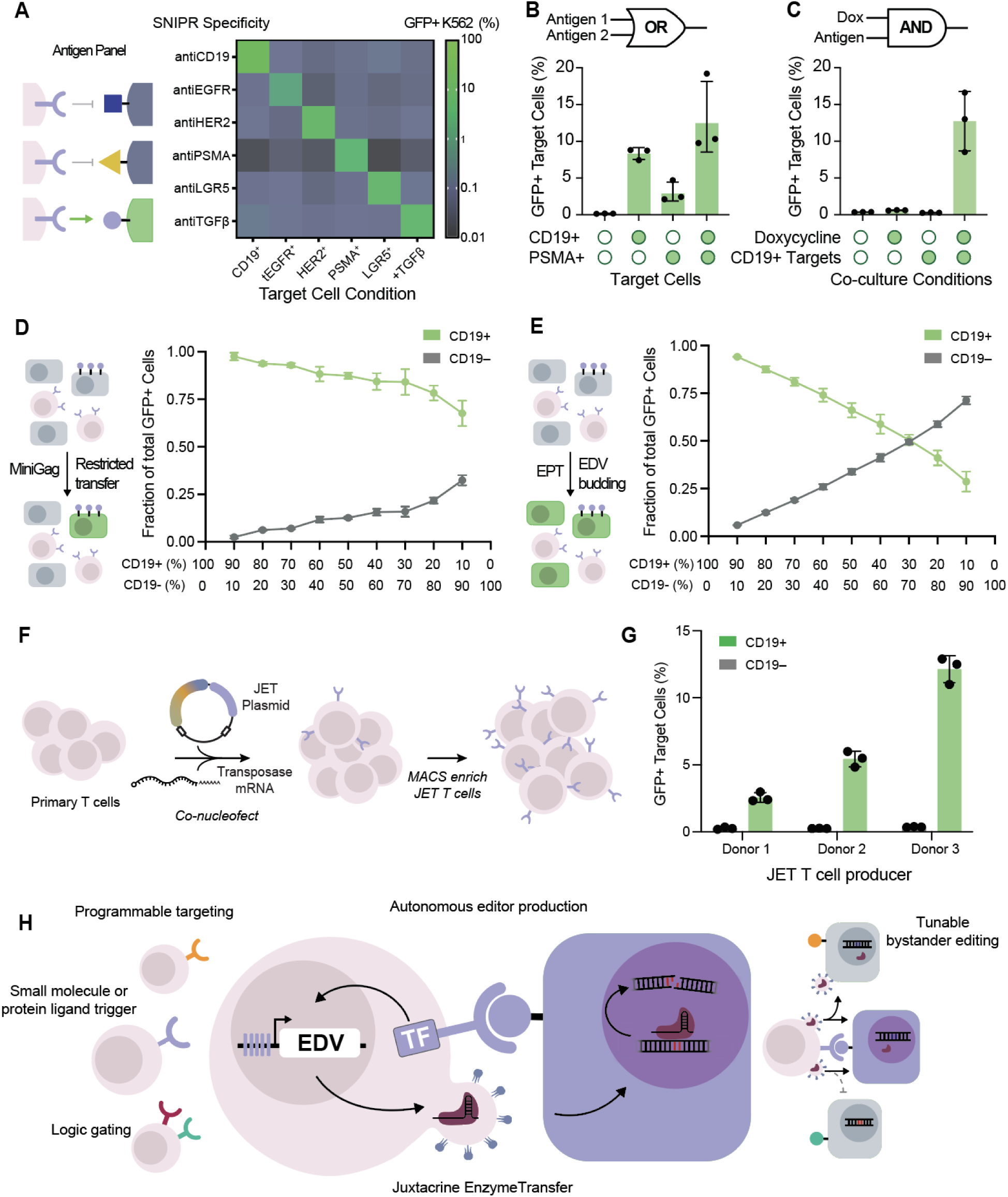
JET T cells enable fully programmable intercellular delivery. A. Combinatorial co-cultures were performed with *lsl*-GFP K562 target cells either transduced with a panel of expression cassettes or in the presence of exogenous TGFβ. (*n* = 3 biological replicates) B. OR-gated delivery of Cre to HEK293T target cells achieved through co-expression of anti-CD19 and anti-PSMA SNIPRs. (*n* = 3 biological replicates) C. Control of the delivery window to HEK293T target cells using an AND–gate achieved through suppression of EDV transcription with TetR. (*n* = 3 biological replicates) D-E. CD19-responsive Jurkat producer cells and target cells are cultured at a 1:1 P:T ratio. The percent of total target cells which express CD19 is varied from 100 to 0. Following 3 days of co-culture, the fraction of total GFP^+^ target cells which are CD19^+^ (“target”) or CD19^−^ (“bystander”) are shown for co-cultures with producer Jurkat cells engineered with either miniGag- (G) or EPT-based (H) Cre-EDV JET circuits. (*n* = 3 biological replicates) F. Schematic for primary JET T cell engineering; primary T cells are nucleofected with Sleeping Beauty transposon and mRNA encoding for the hySB100X transposase. After recovery, JET T cells are enriched by MACS. G. CD19-specific delivery of Cre to HEK293T target cells by primary JET T cells derived from three unique donors. (*n* = 3 technical replicates per unique donor) H. Schematic summarizing JET T cell mechanisms and applications.

Having confirmed that SNIPR modularity extends to programmable EDV production, we next wondered whether delivery specificity could be expanded from a single input^22–24^. Transducing prostate specific membrane antigen (PSMA)-targeted SNIPR producer cells with an additional anti-CD19 SNIPR enabled OR-gated delivery to cells which expressed either CD19, PSMA, or both ligands (Fig. 4B). Similarly, we achieved AND-gated delivery by using two receptors consisting of orthogonal transcription factors (GAL4-VP64 and TetR-VP48) driving separate cassettes encoding the cargo or fusogen, respectively^25,26^ (Fig. S4A, B). Our results confirm that co-expressing multiple synthetic receptors enables Boolean logic-gated delivery to target cells.

Logic gating using two receptors is useful for distinguishing between cell types or tissue microenvironments. We hypothesized that we could additionally achieve both spatial and temporal control of EDV production using a synthetic receptor and a small molecule, yielding cells that only produce EDVs after contact with a target cell only if a drug has also been administered. To engineer such a circuit, we constitutively expressed the Tet repressor protein (TetR) to physically block transcription until doxycycline is administered^27^ (Fig. S4C). Indeed, efficient Cre delivery by these cells was observed only when co-cultured with CD19-expressing target cells in the presence of doxycycline (Fig. 4C).

While complex circuits like those described above can induce EDV production, we hypothesized that EPT-based particles may bud radially from producer cells, rather than unidirectionally towards the target cell. This would imply that neighboring “bystander” cells that do not express the SNIPR-targeted ligand may receive cargo as well, due to the broad tropism of the VSV-G fusogen. In contrast, the miniGag-based constructs, which rely heavily on cell contact for transfer, may show greater target cell specificity. To investigate this, we tested producer cells programmed with either miniGag- or EPT-based EDV cassettes in co-cultures with mixed populations of reporter target and bystander cells (CD19^+^ and CD19^−^, respectively) at a range of target:bystander ratios. Indeed, even when target cells comprised a small fraction of the population, editing by miniGag-EDV producer cells was largely restricted to target cells (Fig. 4D). In contrast, bystander cell editing rose steadily as the proportion of target cells decreased in co-culture with EPT-EDV producer cells, with a majority of edited cells being bystanders at low target:bystander ratios (Fig. 4E). These results collectively demonstrate that delivery by JET Jurkat cells is tunable at the levels of both circuit activation (sensing) and cargo transfer mechanism (output), with different architectures exhibiting different levels of transfer specificity.

### Primary human JET T cells specifically transfer enzymes to target cells

Finally, though Jurkat cells are a useful model for T cell biology, translation to primary T cells is a critical step toward therapeutic application, so we next sought to assess the capacity of primary T cells for cargo production and transfer. To generate primary JET T cells, we co-nucleofected activated primary T cells with mRNA encoding a Sleeping Beauty transposase^28^ and a transposon plasmid donor encoding CD19-responsive Cre EDV circuits^29^. We then isolated primary JET T cells using magnetic-activated cell sorting (MACS) (Fig. 4F; Fig. S4D). Subsequent co-cultures confirmed CD19-specific transfer of Cre by primary T cells from three different donors (Fig. 4G), confirming that the JET circuit can be implemented in primary cells. Critically, unlike CAR based circuits which result in targeted cell death, we observed minimal cytotoxicity to the target cells in our co-cultures (Fig. S4E). This experiment provides an initial demonstration of translating these engineering efforts to a primary T cell chassis.

## Discussion

Here we describe the design and testing of JET T cells engineered to produce and transfer genome editing proteins intercellularly in response to contact with a ligand of interest. While immune cells have many useful traits for cargo delivery, including tissue infiltration and receptor recognition, previous attempts to repurpose lymphocytes for delivery using the innate granzyme-perforin pathway resulted in toxicity to recipient cells^30,31^. We overcame this limitation using an orthogonal circuit that couples antigen sensing to EDV production rather than degranulation, resulting in cells that “seek and deliver” rather than “seek and destroy.” Systemically administered lipid nanoparticles or viral vectors passively bind target cells and are vulnerable to clearance by host cells and serum. In contrast, JET T cells offer the potential to not only perform more complex cellular logic, but to extravasate from the blood to actively infiltrate solid tissues, producing delivery vehicles locally after reaching their intended target.

Over the course of our work, we found that cargo transfer by T-lymphoid cells could occur through two mechanisms depending on the EDV design. Jurkat cells equipped with miniGag-based EDV cassettes primarily deliver editors to cells in direct contact, while Jurkat cells with integrated EPT-based EDV cassettes more efficiently bud particles. These alternative mechanisms provide a unique opportunity to tune the transfer specificity of JET T cells. For extreme specificity in delivery, limiting cargo transfer to cells that stimulate the synthetic receptor would be ideal. Alternatively, certain tissue subpopulations may not have a distinct surface marker but may reside among other cells which do, in which case the ability for cargo to reach neighboring “bystander” cells could be useful (Fig. 4H). Our data suggest that the optimal scaffold choice for particle budding should be determined empirically based on the chosen producer cell chassis and desired output.

SynNotch-equipped T cells have been used to secrete extracellular cargo in response to contact with cells in the brain^32^. We believe JET T cells outline a path for similarly engineering intracellular cargo delivery *in vivo*^33^. Sophisticated logic gating using SNIPRs or other systems could allow greater discrimination of target cell populations, currently difficult to achieve with existing nanoparticle systems. In addition, the inducible production of all xenogenic EDV components may help limit immune clearance until the target tissue has already been reached. We expect this strategy to be compatible with circuits that use alternative synthetic or natural receptors, as well as with alternative producer cell types^34–37^. The repurposing of T cells as couriers for biological cargo establishes a new paradigm in immune cell engineering.

## Limitations of the Study

The biology underlying this cell type-specific difference in mechanism remains unclear, particularly given that both constructs use the same budding motif (HIV-1 p6), and may lie in host factors that specifically respond to retroviral proteins in T cells. Though we were able to select for engineered JET Jurkat cells with antibiotics, the low efficiency of genomic integration for large transgenes like our circuit may limit the large-scale production of primary JET T cells. Additionally, without stimulating T cell activation pathways, JET T cells would not be expected to expand *in vivo* and quiescence may inhibit efficient EDV production. Future engineering efforts may improve production and persistence either through manufacturing optimization or producer cell selection (i.e. CD4^+^ rather than pan-CD3 T cells). Finally, though the immunogenicity of EDV components *in vivo* could be addressed with genome editing (e.g. MHC knockout, CD47 overexpression), VSV-G will necessarily be expressed on the surface of cells after EDV production is induced, which may result in JET T cell clearance by the host immune system. This may provide the advantage of limiting EDV overexpression following contact with the target antigen, but *in vivo* studies will be needed to assess JET T cell persistence.

## Resource Availability

### Lead contact

Further information and requests for resources and reagents should be directed to and will be fulfilled by the lead contact, Jennifer A. Doudna.

### Materials availability

Plasmids generated in this study will be deposited to Addgene upon publication. This study did not generate new unique reagents.

### Data and code availability

Any additional information required to reanalyze the data reported in this paper is available from the lead contact upon request.

## Acknowledgements

We thank Owen Tuck, Peter Yoon, and other members of the Doudna lab for helpful discussions. We also thank Dr. Daniel Humphrys for insightful discussions about the trimeric 1WA3 nanocage mutant. K.M.W. is a recipient of the Bakar BioEnginuity Impact Grant. J.A.D. is an investigator of the Howard Hughes Medical Institute (HHMI). This work was supported by the NIH/NIAID HIV Accessory & Regulatory Complexes (HARC) Center grant U54 AI170792. J.A.D. also receives support from National Institutes of Health (U19AI135990, UH3AI150552, U19NS132303, and R21HL173710), National Science Foundation (2334028), Department of Energy (DE-AC02-05CH11231, 2553571, and B656358), Lawrence Livermore National Laboratory, Apple Tree Partners (24180), UCB-Hampton University Summer Program, Mr. Li Ka Shing, Koret-Berkeley-TAU, Emerson Collective, and the Innovative Genomics Institute.

## Author Contributions

Conceptualization, K.M.W., and J.A.D.; experimental studies, K.M.W., M.M., K.C., E.M., R.P., E.C., and D.C.; data analysis, K.M.W., M.M., and W.N.; supervision, J.B. and J.A.D.; and manuscript writing, K.M.W. and J.A.D. with critical input from W.N., M.M. and J.B..

## Materials and Methods

### Plasmid Construction & Preparation

All plasmids were cloned in-house using ligation cloning, Gibson assembly, or Golden Gate assembly. All oligonucleotides were ordered from IDT. PCR products were generated with Q5 High-Fidelity 2X PCR master mix. Coding sequences for SNIPR and EDV components were codon-optimized for human T cells using the GenSmart Codon Optimization online tool and ordered as DNA fragments from IDT or Twist Bioscience. Sequences of all plasmids were verified using Plasmidsaurus, Quintara, or Azenta. Plasmids used in this study will be listed in the Key Resources Table. scFv sequences were derived from published patents and scientific literature. Plasmids for nucleofection were purified using a Qiagen QIAprep Spin Miniprep Kit or ZymoPure II Plasmid Maxiprep Kit.

### Cell Culture

Jurkat, HEK293T, Lenti-X 293T, K562, and A549 cell lines were obtained from the UC Berkeley Cell Culture Facility. NALM6 cells were obtained from ATCC. Jurkat, K562, and NALM6 cells were maintained in RPMI-1640 (Gibco) with 10% FBS (VWR) and 100U/ml penicillin/streptomycin (Gibco). HEK293T, Lenti-X 293T, and A549 cells were maintained in DMEM (Corning) with 10% FBS and 100U/ml penicillin/streptomycin. NPCs were isolated from embryonic day 13.5 Ai9-tdtomato homozygous mouse brains. Cells were cultured as non-adherent neurospheres in NPC medium (DMEM/F12 with GlutaMAX supplement, sodium pyruvate, 10mM HEPES, 1X nonessential amino acids, 100U/ml penicillin/streptomycin, 55µM 2-mercaptoethanol, B-27 and N-2 supplements, 20 ng/ml bFGF and 20 ng/ml EGF. NPCs were passaged using the MACS Neural Dissociation Kit according to the manufacturer’s instructions.

### Co-culture Assays

Co-cultures were performed in 96-well plates. 50,000 target cells were seeded into each well alone or with 50,000 producer cells in 150µl total complete RPMI. Where applicable, doxycycline was added to a final concentration of 500 ng/ml. For neural progenitor cell co-cultures, three days prior to co-cultures and supernatant treatments, 5,000 NPCs were seeded in complete NPC medium per well of a 96-well plate pre-coated with a solution containing poly-DL-ornithine hydrobromide, laminin, and fibronectin bovine plasma. On the day of the assay, four wells were trypsinized and counted to ensure a true 1:1 producer:target ratio. Producer cells were added in complete NPC medium.

For Cre transfer assays, flow cytometry analysis was performed after 72 hours of co-culture. For Cas9 transfer assays, to allow for β2M protein reduction, producer and target cells were passaged together after 72 hours and split 1:5 into a new 96-well plate. Flow cytometry analysis was then performed after another 72 hours (day 6).

For primary organoid co-cultures, 2e4 dox inducible Cre-EDV jurkat cells and 1e4 cells from trypsinized gastric, small intestine or colon organoids were mixed and seeded together in Matrigel droplets. Doxycycline added at 500ng/mL either immediately or 24 hours after seeding. At endpoint, hoechst 34580 (ThermoFisher) was added at a final concentration of 5ug/ml, incubated for an hour at 37°C, and imaged via fluorescence confocal microscopy as well as bright field imaging.

For CD19+ cancer organoids, 1e4 hCD19-targetting jurkats and 2e3 trypsinized *Kras*^G12D^;*Trp53*^-/-^; hCD19-shAPC; LSL-tdTomato organoids were mixed and seeded together in Matrigel droplets. At endpoint, hoechst 34580 was added at a final concentration of 5ug/ml, incubated for an hour at 37°C, and imaged via fluorescence confocal microscopy as well as bright field imaging.

### Supernatant Transfer & Transwell Budding Assays

Transwell assays were performed exclusively with producer Jurkat cells in the bottom chamber to avoid the possibility of migration through the porous insert. In a HTS Transwell 96-Well Plate with 3.0µm pores, 200,000 Jurkat producer cells were plated in the bottom chamber with 25,000 HEK293T target cells in the top chamber. Final doxycycline concentration was either 0 or 500 ng/ml. After 6 days of co-culture, HEK293T cells in the top chamber were trypsinized and analyzed for β2M surface expression by flow cytometry.

For supernatant transfer assays, producer Jurkat cells were cultured at 6.66e5 cells per milliliter in serum-free OptiMEM (for HEK293T targets) or complete NPC media (for NPC targets) in the presence of 500 ng/ml doxycycline for 36-48 hours. Cultures were then centrifuged at 500g for 5 minutes and supernatant was passed through a 0.45µm filter before adding to HEK293T or Ai9 target cells. Target cells were then passaged after 72 hours, at which point tdTomato expression was assessed by flow cytometry for NPC targets, or replated with final readout of β2M knockout after three more days (day 6) by flow cytometry for HEK293T targets.

### Flow Cytometry

During co-culture passage at day 3 or assay endpoints at day 3 or day 6, supernatant was carefully transferred to a new 96-well plate before 35µl of 0.25 % trypsin-EDTA was added to the adherent cell monolayer. To quench trypsin, spent supernatant was added back to ensure non-adherent Jurkat cells were maintained. Cells were then centrifuged at 300g for 5 minutes, washed once with PBS/BSA, resuspended in 50µl staining master mix, and incubated at 4°C for 30 minutes. Excess antibody was then washed out and cells were analyzed on an Attune NxT flow cytometer with autosampler. For primary T cell co-cultures, prior to antibody staining, cells were labeled with Zombie NIR dye in PBS to enable viability measurement.

### Primary T Cell Culture

Peripheral blood mononuclear cells were purchased from ATCC. T cells were isolated using an EasySep™ Human T Cell Isolation Kit (STEMCELL) according to the manufacturer’s instructions and rested overnight in complete T cell media (X-VIVO 15 with 5% FBS, 10mM N-acetyl-L-cysteine, and 50µM 2-Mercaptoethanol) without cytokines. The following day, cells were activated with DynaBeads at a 1:1 bead:cell ratio at a density of 1e6 cells/ml in the presence of 500U/ml IL-2, 5ng/ml IL-7, and 5ng/ml IL-15. After 48 hours of stimulation, cells were debeaded and either immediately prepared for nucleofection or passaged in complete T cell media with 100U/ml IL-2. T cells post-activation were cultured every other day at a density of 1e6 cells/ml with fresh IL-2 addition.

### Magnetic-Activated Cell Sorting

Primary JET T cells were isolated using an EasySep™ Release Human PE Positive Selection Kit (STEMCELL) according to the manufacturer’s instructions. Briefly, cells were resuspended at 1e8/ml in EasySep Buffer (STEMCELL). Cells were treated with FcBlock and antiHA-PE (BioLegend) and isolated using an EasyEights EasySep magnet. Cells were eluted from beads with 1x release buffer, centrifuged and resuspended in complete T cell media.

### Organoid Generation & Culture

Non-cancerous gastric, colon, and small intestine organoids were derived from Ai9 mice and cultured with WNT3a, RSPO3, noggin (WRN) conditioned media supplementation to Advanced DMEM/F-12^38^. Media was supplemented with 50ng/ml mouse EGF.

Colon cancer organoids were generated from a *Kras*^LSL-G12D^;*Trp53*^flox/flox^ mouse on a C57Bl/6 background. Cells were transfected with a non-integrative Cre-containing plasmid to generate *Kras*^G12D^;*Trp53*^-/-^ (KP) organoids and were selected with 10uM nutlin-3 and 1uM gefitinib for two weeks. Subsequently, KP organoids were transfected with an integrative piggyBac plasmid expressing EF1a::human CD19 with an shAPC in 3’ UTR and were selected via WRN withdrawal^39^. Finally, these *Kras*^G12D^;*Trp53*^-/-^; hCD19-shAPC organoids were transfected with a piggyBac plasmid containing CAG>LSL-tdTomato and selected with blasticidin. This *Kras*^G12D^;*Trp53*^-/-^; hCD19-shAPC; LSL-tdTomato colorectal cancer organoid reporter line was maintained in advanced DMEM/F-12, 5% fetal bovine serum, 2 mM GlutaMAX, and 100 U/mL penicillin-streptomycin.

Both Ai9 gastrointestinal tract organoids and colorectal cancer organoids were cultured in 9ul domes and seeded in 48-well tissue culture treated plates. Plates were placed in a humidified 37C incubator, upside down, and allowed to solidify for 15-25 minutes. For all experiments, Corning growth factor reduced Matrigel was used at 75% v/v. For all experiments, growth media was added at a volume of 300–600ul.

### Microscopy

Neural progenitor cell co-cultures were imaged in 96-well plates on an EVOS FL inverted fluorescence microscope, equipped with an LPlanFL PH2 10x/0.30NA objective lens, an RFP 2.0 (542/20 ex, 593/40 em) light cube, and a Sony ICX445 monochrome CCD camera (1280 x 960 pixel resolution, 1.3 Megapixels).

The organoids were imaged longitudinally on an ImageXpress Confocal HT.ai High-Content Imaging System. Images were acquired at 4× magnification using the DAPI and Texas Red lasers, as well as the TL50 Brightfield acquisition mode. One field of view was acquired per well, with 10 z-slices.

To quantify fluorescent organoid images, maximum intensity projection images were generated from the10 z-planes captured for each well. A background subtraction was performed on the projection images and then total tdTomato fluorescence was quantified by integrating the intensity values of every pixel.

### Lentivirus Production

3.8 million Lenti-X HEK293T cells were seeded into a 10cm tissue culture-treated plate in 10ml complete media. The next day, transfection plasmids (1µg pMD2.G, 10µg psPAX2, 10µg lentiviral transgene) were added to 400µl Opti-MEM. 63µl PEI (concentration) was added to the DNA solution, vortexed briefly, and incubated at room temperature for 15-30 minutes before adding the transfection mix dropwise to the cells. Media was aspirated and replaced with 10ml pre-warmed serum-free OptiMEM the next morning. Lentivirus production was left to occur for two days, at which point supernatant was harvested, purified through a 0.45µm filter, and either added to cells directly or concentrated 20-fold using Lenti-X Concentrator according to the manufacturer’s instructions.

### Lentivirus Transductions

1e5 cells were seeded into wells of a 24-well plate. Cells were transduced in 50% complete media, 50% Opti-MEM with several doses of concentrated lentivirus (0.5, 5, or 50 µl of 20X concentrated lentivirus). Cells received a full media change 48 hours later with complete media. For *loxP*-GFP reporter cell lines, cells were then selected with 1µg/ml puromycin. For logic-gated producer cell lines, cells were selected with 8µg/ml blasticidin. The lowest dose of virus that resulted in sufficient resistant cells was expanded and used for further assays. For target cells engineered to stably express a given antigen, cells were sorted to enrich for successfully transduced ligand-positive cells. For parental reporter cells, GFP-negative cells were also sorted to remove prematurely GFP+ cells, which appeared likely due to template switching during lentivirus reverse transcription.

### In vitro transcription of mRNA

*In vitro* transcription of mRNA was performed using the HiScribe T7 ARCA mRNA Kit (with Tailing) according to the manufacturer’s instructions. Briefly, the T7-hySB100X plasmid was linearized by digestion with XhoI at 37°C for 5 hours and purified using a QIAquick PCR Purification Kit. 1µg DNA template was incubated with ARCA/NTP mix, DTT, and T7 RNA polymerase at 37°C for at least one hour. Reaction was treated with 2µl DNase I at 37°C for 15 minutes. mRNA was then poly(A)-tailed by incubation with *E. coli* Poly(A) Polymerase at 37°C for 45 minutes. mRNA was purified by lithium chloride precipitation.

### Nucleofection

Nucleofections were performed using a Lonza 4D-Nucleofector 96-well Unit according to the manufacturer’s instructions. Jurkat cells were seeded at 1e5 cells/ml three days prior to nucleofection. 2e5 cells were centrifuged at 92g for 10 minutes and resuspended in 20µl of SE Buffer. Cells were mixed with 800ng of transposon plasmid and 200ng of transposase plasmid, transferred to the nucleofection cuvette, and electroporated using pulse code CL-120. Immediately after nucleofection, 80µl pre-warmed media was added to each cuvette well and cells were incubated 10-15 minutes at 37°C. Cells were then moved to wells of a 96-well plate containing 50µl pre-warmed media.

Following activation, primary T cells were removed from activation beads, counted, and centrifuged at 92g for 10 minutes. T cells were resuspended in P3 Buffer at a concentration of 2e6 cells per 20µl, mixed with 2.5µg transposon plasmid DNA and 0.5µg hySB100X mRNA, and transferred to nucleofection cuvettes. Cells were electroporated using pulse code EO-151. Immediately after nucleofection, 80µl pre-warmed media was added to each cuvette well and cells were incubated 10-15 minutes at 37°C. Cells were then moved to wells of a 96-well U-bottom plate containing 200µl pre-warmed media with IL-2 and rested overnight. The next day, cells were passaged at 1e6 cells/ml with IL-2.

### Western Blot

Jurkat cells were cultured in the presence of doxycycline for 48 hours. Cells were then pelleted, washed once with cold PBS, resuspended in Pierce RIPA Buffer containing Halt Protease Inhibitor, and lysed on ice for 15 minutes. Lysates were pelleted at 20,000xg at 4°C to remove insoluble debris, and supernatant was transferred and quantified using a Pierce BCA Assay Kit. 5 µg protein lysate was denatured at 95°C for 4 minutes and run on a 10% SDS-PAGE gel for 1 hour at 150 V. Proteins were transferred to a PVDF membrane at 90 V at 4°C for 1 hour in transfer buffer consisting of 25 mM Tris, 192 mM glycine, and 20% (v/v) methanol. Membranes were blocked for 1 hour at room temperature on an orbital shaker in blocking buffer consisting of 1× PBS with 0.1% Tween-20 (PBS-T) with 5% non-fat milk. Membranes were washed four times with 1□PBS containing 0.1% Tween-20 (PBS-T), then incubated while rocking overnight at 4°C in blocking buffer with anti-GAPDH (1:1000) and either anti-FLAG (1:2,000) or anti-VSV-G (1:1,000) antibodies. Membranes were washed four times with PBS-T, then incubated with secondary antibody (1:20,000) while rocking for 1 hour at room temperature. The membranes were then washed four times with PBS-T, and imaged on a LICOR imager (LICORBio).

---

**Supplementary Figure 1:**
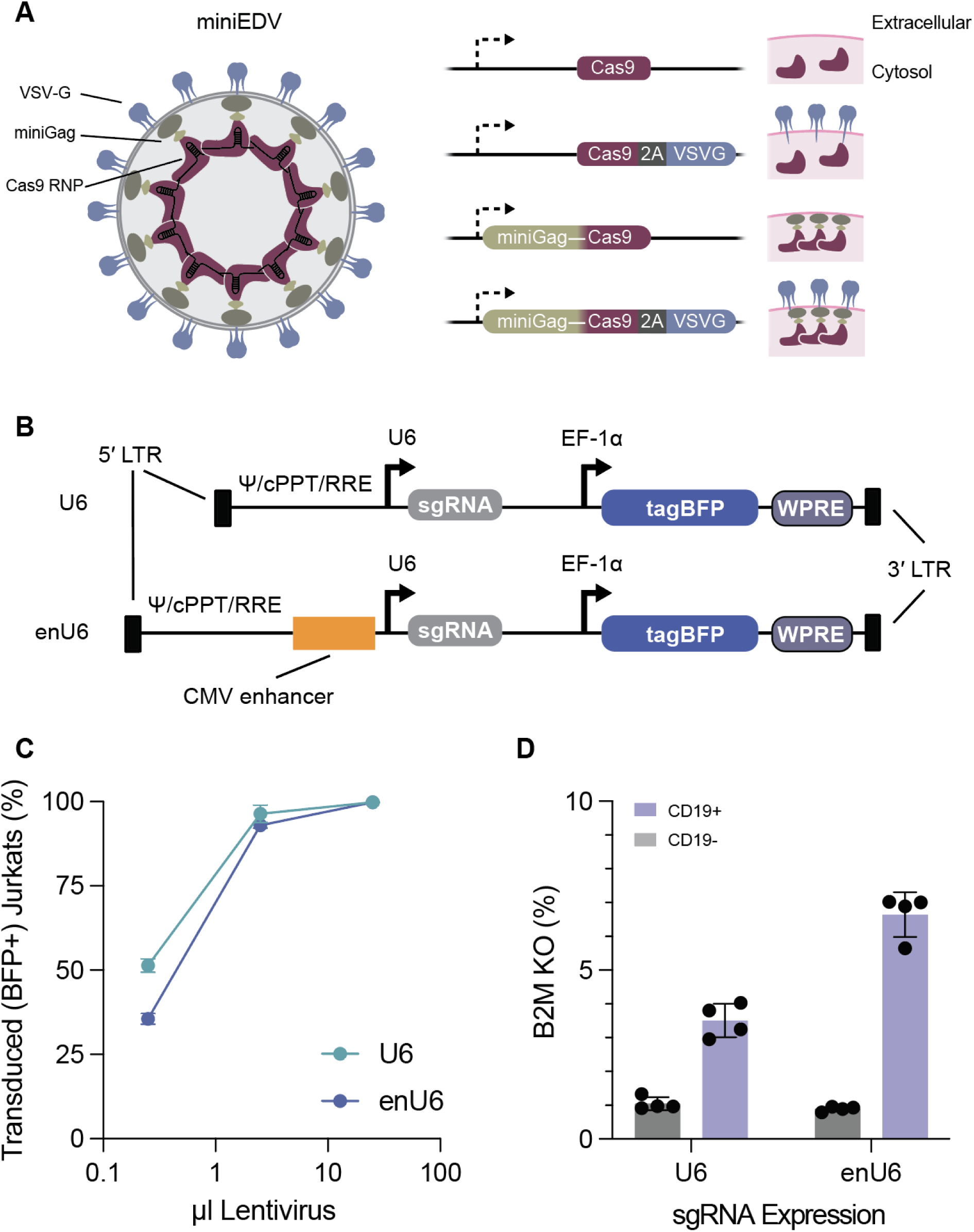
A. Cartoon of miniGag EDV structure showing VSV-G fusogen (blue), miniGag (green), and Cas9 RNP (red). Right: Constructs used in Figure 1 with expected outcomes in producer cells at the plasmid membrane. B. Schematic of lentiviral vectors used to install *B2M*-targeted sgRNA expression in producer and target cells, using either U6 alone (top) or U6 downstream of the CMV enhancer (bottom). C. Transduction (BFP^+^) in Jurkat producer cells quantified by flow cytometry following volume-based transductions with either of the constructs in (B). (*n* = 4 biological replicates) D. *B2M* editing in target cells following co-culture with Jurkat producer cells transduced with the 50µl dose of lentivirus from (C). (*n* = 4 biological replicates)

**Supplementary Figure 2:**
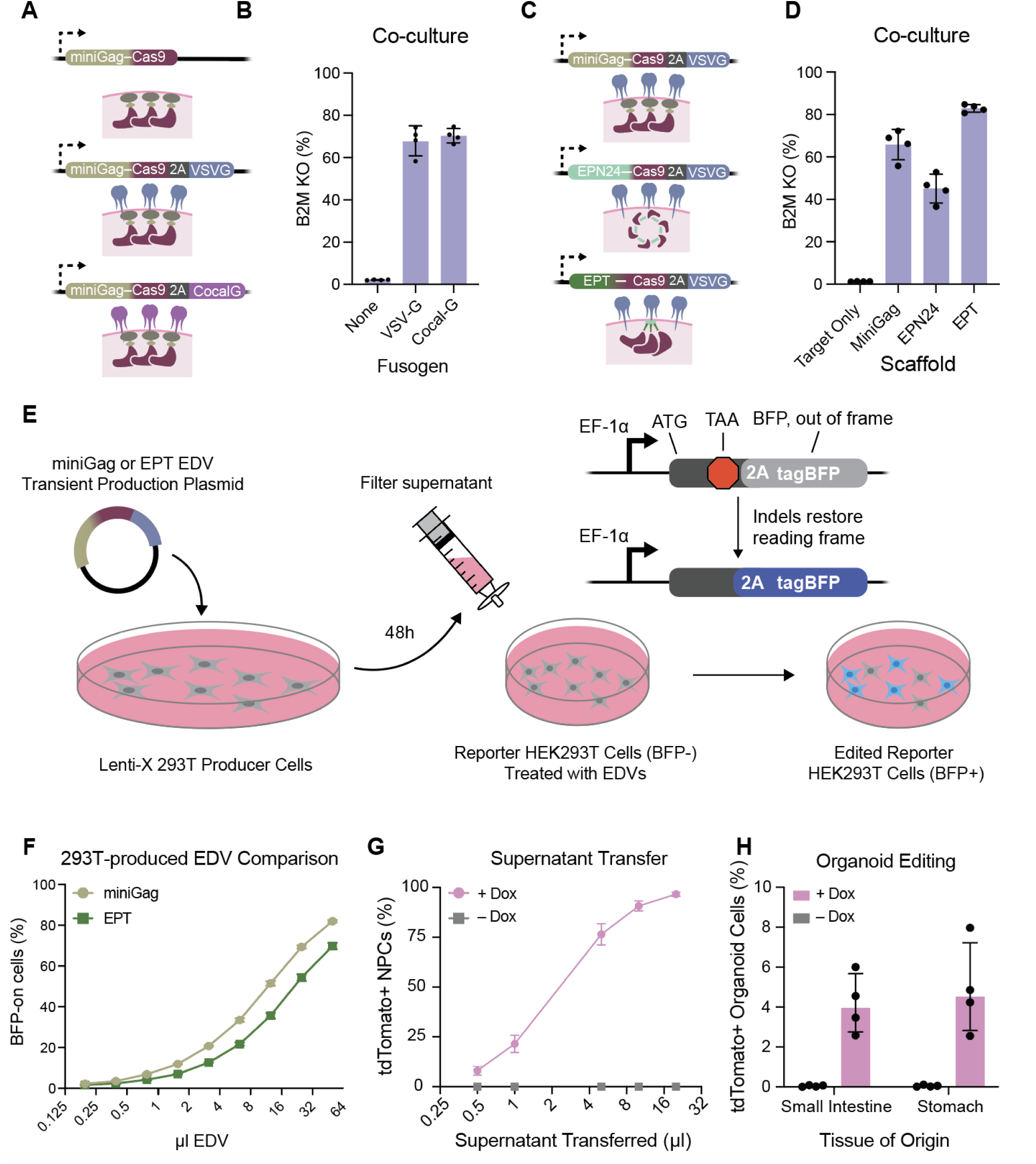
A. Visual representations of EDV cassettes encoding for no fusogen, VSV-G, or Cocal-G. B. Equivalent target cell editing using Jurkats expressing miniGag-Cas9 EDVs pseudotyped with VSV-G or Cocal-G measured by flow cytometry following 6 days of co-culture. (*n* = 4 biological replicates) C. MiniGag, EPN24, and EPT EDV cassette visual representations. D. Extension of Figure 2B including EPN24; Loss of β2M measured by flow cytometry indicates EPN24-based EDV construct leads to less efficient Cas9 transfer following co-culture in the presence of doxycycline. (*n* = 4 biological replicates) E. Schematic of Cas9 editing reporter assay. Plasmids encoding miniGag- or EPT-based EDVs driven by the CAG promoter and a U6-sgRNA cassette were transfected into Lenti-X 293T cells. EDVs were harvested and filtered from supernatant and used to treat HEK293T targets with genomically integrated indel reporter cassettes, in which a premature stop codon prevents BFP expression until Cas9-mediated indels restore the BFP reading frame. Editing is quantified as BFP expression by flow cytometry. F. Comparison of editing results after treatment with miniGag or EPT- based Cas9 EDVs produced by Lenti-X 293T cells. (*n* = 3 biological replicates) G. Jurkat cells were cultured in doxycycline to produce EPT-Cre EDVs. Supernatant was filtered and added to Ai9 NPCs. tdTomato fluorescence in NPCs indicates successful delivery of Cre. (*n* = 4 biological replicates) H. Flow cytometry quantification of Cre delivery to organoids from Ai9 mouse stomach and small intestines following co-culture with producer Jurkats. (*n* = 4 biological replicates)

**Supplementary Figure 3:**
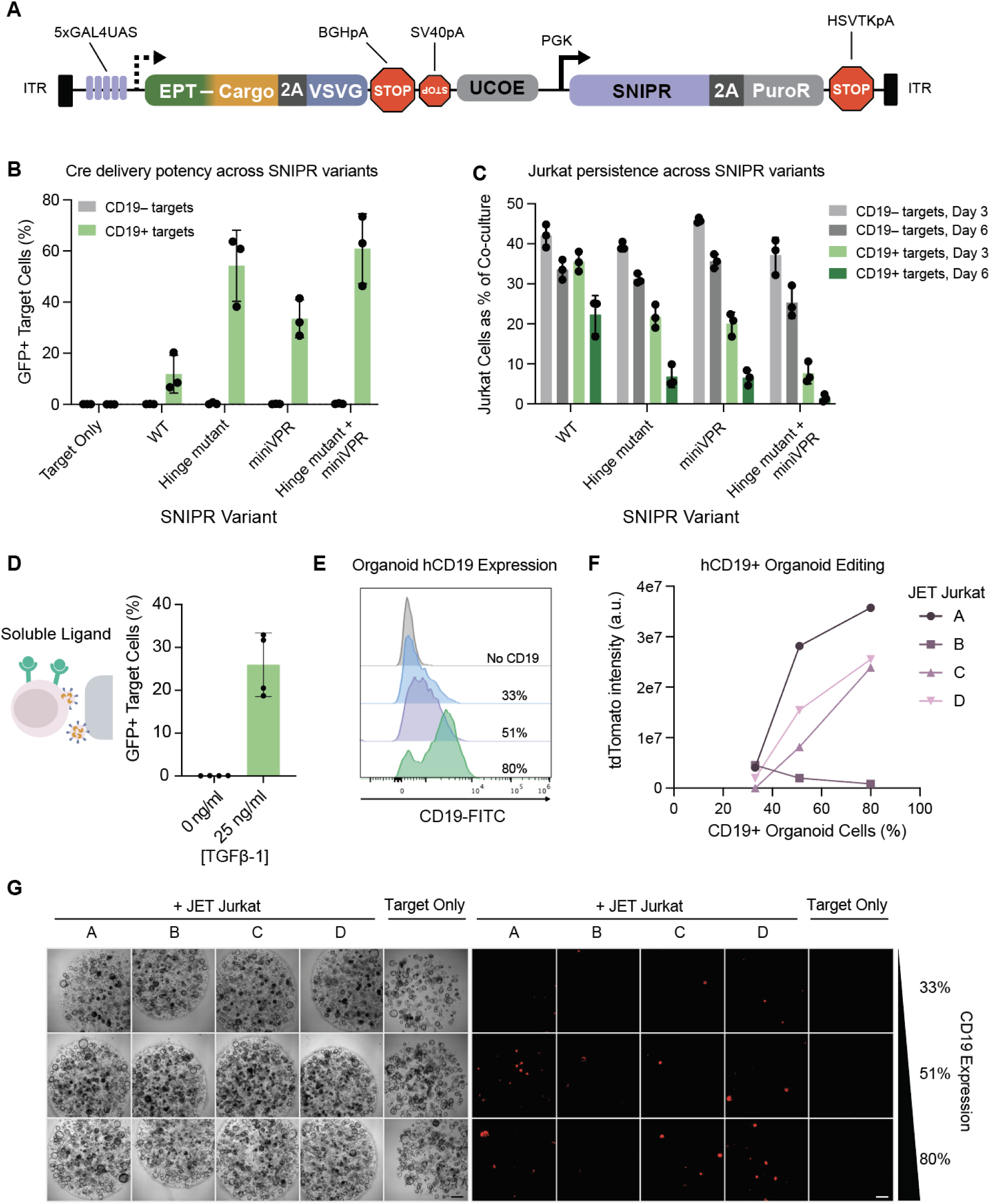
A. Map of all-in-one JET Sleeping Beauty construct. 5xGAL4UAS + minimal promoter drives inducible EDV expression. SNIPR and puromycin resistance are driven by a PGK promoter with ubiquitous chromatin opening element (UCOE). As the UCOE is essentially a bidirectional promoter, an inverted SV40 poly(A) sequence was included to terminate antisense transcription^40^. B. JET Jurkats were generated with previously described SNIPR variants. GFP fluorescence in target cells after 3 days of co-culture is shown for each condition. (P:T = 1, *n* = 3 biological replicates) C. Quantification of Jurkats as a percent of total cells in co-culture after 3 and 6 days after initially being seeded at 50%, using the SNIPR variants tested in (B). (*n* = 3 biological replicates) D. Cre delivery assessed by flow cytometry measurement of GFP expression in target *lsl*-GFP HEK293T cells following 3 days of co-culture with JET Jurkat cells (anti-TGFβ) in the absence or presence of 25 ng/ml TGFβ1. (*n* = 4 biological replicates) E. Human CD19 expression on engineered shAPC, *Kras*^G12D^*;Trp53*^-/-^ mutant (AKP) colon cancer organoid lines. F. Quantification of fluorescent intensities from co-cultures with various JET Jurkat producers with organoids expressing hCD19 at various levels. Each point represents the average of technical replicates for a given Jurkat biological replicate. Lack of function in JET Jurkat line B is likely due to transgene silencing, resulting in reduction in transfer efficiency as engineered producer cells are kept in culture over time^41^. G. Brightfield (left) and red fluorescence (right) representative images of *lsl*-tdTomato hCD19+ organoids after four days of co-culture with CD19-responsive JET Jurkat cells, related to (F). (scale bar = 500 µm)

**Supplementary Figure 4:**
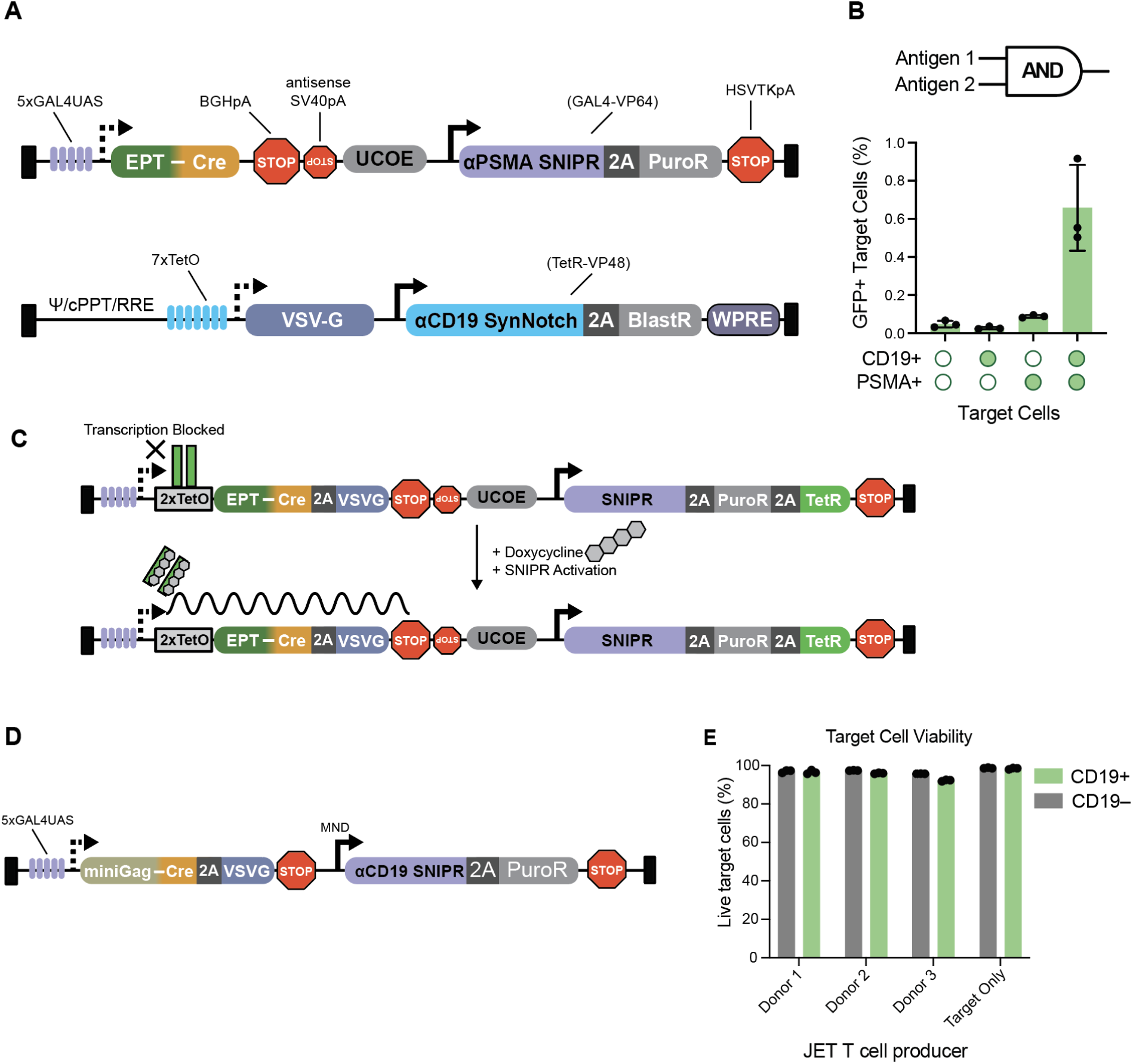
A. Schematics of dual-ligand AND-gated constructs. B. Flow cytometry measurement of GFP expression in target HEK293T cells expressing CD19, PSMA, both, or neither following 3 days of co-culture with AND-gated JET Jurkat cells. C. Schematics of ligand+doxycycline AND gated construct. (Related to Fig. 4C) D. Schematic of Sleeping Beauty construct used for primary JET T cell generation. E. Viability data for target cells following 3 days of co-culture with primary T cells. (Related to Figure 4G)

## Declaration of generative AI and AI-assisted technologies in the manuscript preparation process

During the preparation of this work the authors used Claude in order to improve readability. After using this tool, the authors reviewed and edited the content as needed and take full responsibility for the content of the published article.

